# Challenging the Astral^™^ mass analyzer - up to 5300 proteins per single-cell at unseen quantitative accuracy to study cellular heterogeneity

**DOI:** 10.1101/2024.02.01.578358

**Authors:** Julia A. Bubis, Tabiwang N. Arrey, Eugen Damoc, Bernard Delanghe, Jana Slovakova, Theresa M. Sommer, Harunobu Kagawa, Peter Pichler, Nicolas Rivron, Karl Mechtler, Manuel Matzinger

**Author notes:** These authors contributed equally (Julia A. Bubis, Manuel Matzinger).

## Abstract

A detailed proteome map is crucial for understanding molecular pathways and protein functions. Despite significant advancements in sample preparation, instrumentation, and data analysis, single-cell proteomics is currently limited by proteomic depth and quantitative performance. We combine a zero dead-end volume chromatographic column running at high throughput with the Thermo Scientific™ Orbitrap™ Astral™ mass spectrometer running in DIA mode. We demonstrate unprecedented depth of proteome coverage as well as accuracy and precision for quantification of ultra-low input amounts. Using a tailored library, we identify up to 7400 protein groups from as little as 250 pg HeLa at a throughput of 50 samples per day (SPD). We benchmark multiple data analysis strategies, estimate their influence on FDR and show that FDR on protein level can easily be maintained at 1 %. Using a two-proteome mix, we check for optimal parameters of quantification and show that fold change differences of 2 can still be successfully determined at single-cell level inputs. Eventually, we apply our workflow to A549 cells yielding a proteome coverage of up to 5300 protein groups from a single cell, which allows the observation of heterogeneity in a cellular population and studying dependencies between cell size and cell-cycle phase. Additionally, our workflow enables us to distinguish between *in vitro* analogs of two human blastocyst lineages: naïve human pluripotent stem cells (epiblast) and trophectoderm (TE)-like cells. Gene Ontology analysis of enriched proteins in TE-like cells harmoniously aligns with transcriptomic data, indicating that single-cell proteomics possesses the capability to identify biologically relevant differences between these two lineages within the blastocyst.

## INTRODUCTION

Single-cell proteomics (SCP) by mass spectrometry has evolved to a powerful technique to investigate cellular heterogeneity with increasing coverage and throughput. To tackle the challenge of ultra-low input amounts, a multitude of miniaturized and automatized sample preparation workflows have been developed by several labs.^1–8^ The success of SCP, however, is also tightly connected to technological improvements in high performance, nano-flow, liquid chromatography (LC) as well as to high resolution, high sensitivity mass spectrometry (MS). The SCP field initiated its success by using isobaric labels for multiplexing to improve sensitivity and throughput.^1^ With more sensitive and fast mass spectrometers being commercialized, now more and more groups focus on label-free workflows^9^ and employ data independent acquisition (DIA) workflows combined with short LC gradients. With these DIA workflows optimal reproducibility across runs is achieved, while maintaining throughput at acceptable levels ranging from 40 to up to 180 samples per day (SPD).^10–12^ Improvements in data analysis algorithms allows for confident identification of very low abundant peptides hidden within highly chimeric spectra.

The recent introduction of the Thermo Scientific™ Orbitrap™ Astral™ mass spectrometer^13^ further extends the reachable sensitivity and acquisition speed by combining the established Thermo Scientific™ Orbitrap™ analyzer with the newly developed Thermo Scientific™ Astral™ analyzer, in a single instrument. The Astral analyzer combines high speed (up to 200 Hz) at high resolution and sensitivity, with nearly lossless transmission and a high dynamic range.^13^ The instrument allows for high acquisition speed by simultaneously utilizing the Orbitrap analyzer for MS1 spectra and the Astral analyzer for MS2 spectra at a resolution sufficient to even resolve Tandem Mass Tag (TMT) reporter ions and at a sensitivity superior to the Orbitrap analyzer.^13^

Here we investigate the performance of the Orbitrap Astral MS in combination with the Aurora™ Ultimate TS 25 cm nanoflow Ultra High-Performance Liquid Chromatography (UHPLC) column that comes with a fully integrated source interface to minimize peak broadening and yield the highest possible signal intensity. We optimize the gradient length to yield maximum proteome coverage and assess the accuracy and precision of quantification.

Capitalizing on the synergistic effects of the aforementioned methodological improvements, SCP now enters a stage where it serves as an important tool for biologists to complement single-cell genomics data.^14^ We apply our improved workflow to investigate the cellular heterogeneity of A549 human lung cancer cells. Additionally, we employed optimized SCP on naïve human pluripotent stem cells (hPSC), which recapitulate the pre-implantation EPI and on TE-like cells differentiated from naïve hPSCs. During the in vitro fertilization (IVF) process, only about 40% of fertilized eggs are estimated to reach the blastocyst stage with sufficient quality for transfer to the mother’s uterus.^15^ The criterion for selecting blastocysts largely depends on morphological characteristics, such as the expansion of the outer trophectoderm (TE) and a well-clustered inner epiblast (EPI) population.^16,17^ However, our understanding of the mechanisms regulating blastocyst morphology remains limited. This limitation is primarily due to restricted access to human embryos and the lack of technologies capable of low-input detection methods. In this context, high-sensitivity SCP emerges as a powerful tool. It holds promise for unraveling the mechanisms underlying the regulation and functionalization of blastocyst morphology, including aspects like TE epithelialization, cavity formation, and tissue segregation between the EPI and TE.

## RESULTS

### Combination of zero dead volume chromatography with the Orbitrap Astral MS, DIA and a tailored library results in unprecedented proteomic coverage

We combined the Orbitrap Astral MS with the Thermo Scientific™ FAIMS Pro interface and the Auroa Ultimate 25 cm TS column with an integrated emitter tip and zero dead volume, aiming to maximize sensitivity for ultra-low input proteomic samples. To have a representative standard of comparable input to a single cell for evaluation and benchmarking, we decided to inject 250 pg of commercial HeLa and K562 peptides. The bulk digests were diluted in 0.1% TFA containing 0.015 % N-Dodecyl-β-D-maltoside (DDM) to improve peptide solubility; reproducibility and quantitation accuracy.^18^ Gradient lengths were varied from 14 to 38 min (Supplemental Table 1), which corresponds to reachable throughputs of 30 – 80 SPD, including sample loading as well as column equilibration, and washing time. During loading and washing flow rates were increased to improve throughput and lower flow rates were applied during peptide elution to enhance sensitivity.^12,19,20^ As shown in Figure 1, we found a sweet spot around 50 SPD yielding maximal identifications, both on precursor and protein group levels for both cell lines. Of note, the data was analyzed library-free. When replicates were searched together, the number of protein groups identified was boosted by 6.3 – 10.9 % and 4.5 – 7.2 % for HeLa and K562 respectively. In addition, by allowing for matching, replicate measurements nicely align, leading to a data completeness close to 100 % and vanishing error bars for ID numbers compared to the method evaluation mode (Figure 1).

**Figure 1:**
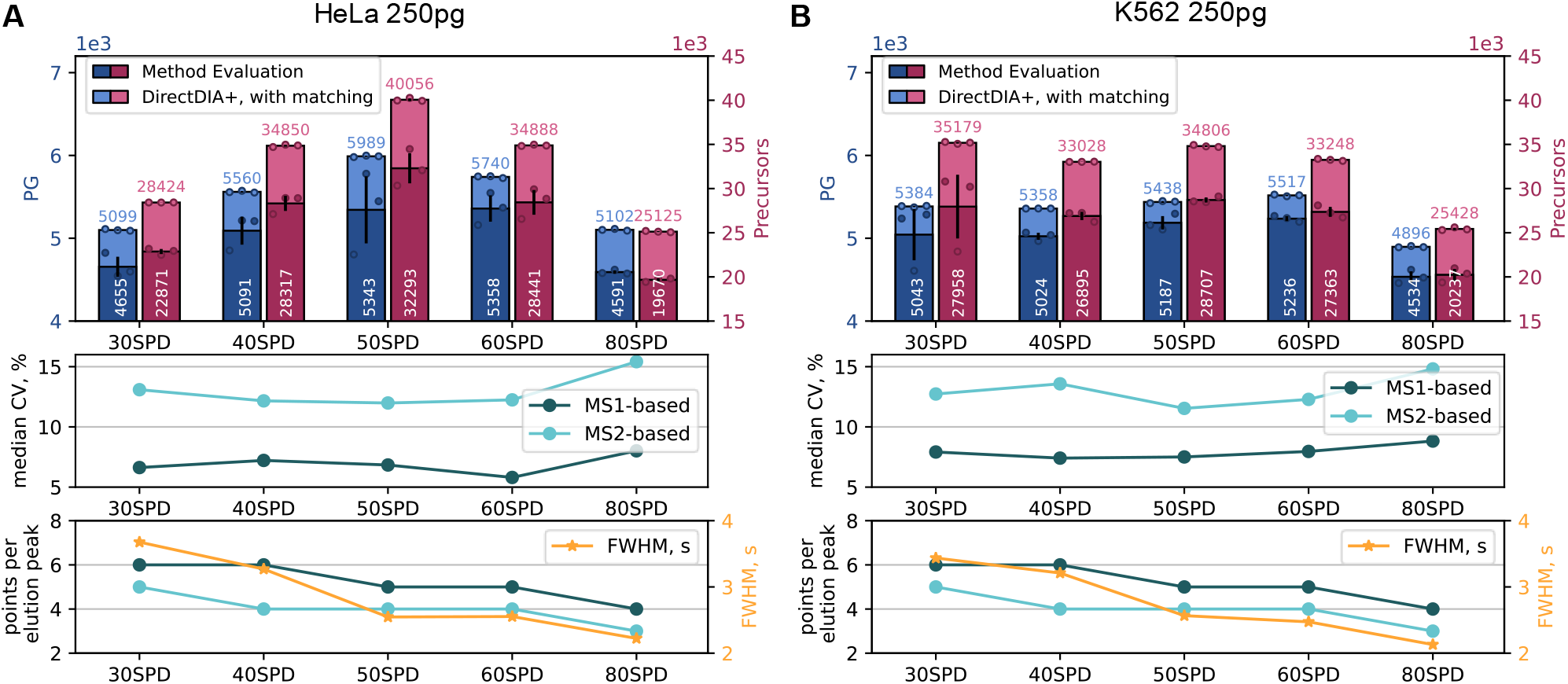
Finding an optimal gradient length to maximize protein IDs and maintain data quality. 250 pg of HeLa (**A**) or K562 (**B**) peptides from diluted bulk digest were injected each. Peptides were separated using a gradient of altered length ranging from throughputs of 30 – 80 SPD. Gradient details are depicted in Supplemental Table 1. Data was recorded in DIA mode on the Orbitrap Astral MS and analyzed in direct DIA+ mode using Spectronaut 18. Top graphs: Circles indicate identified protein groups (PG) or precursors at 1% FDR in individual replicates, bars indicate their means, while error bars indicate standard deviations with n=3. Dark colors represent identified proteins in method evaluation mode, with no matching across replicates and light colors indicate boosted ID numbers when allowing for matching. Middle graphs: The dots indicate the obtained median CV on protein level for each gradient length (DirectDIA+ with matching) when performing quantification on MS1 or MS2 level respectively. Bottom graphs: The dots indicate the median number of datapoints per precursor that were used for quantification for each gradient length on MS1 or MS2 level respectively as well as the median FWHM of elution peaks.

Based on our results, we hypothesize that shorter gradients are advantageous for ultra-low input samples as they produce sharper, and hence more intense, chromatographic elution peaks. Once a maximum is reached at 50 SPD, faster gradients potentially suffer from insufficient separation power and greater spectrum complexity, which adversely affects identification numbers. Fast gradients with very narrow elution peaks further reduce the number of datapoints per peak (Figure 1, bottom graphs). Although this had no strong impact on the coefficient of variation (Figure 1, middle graphs), it seems obvious that a higher number of datapoints per peak results in better peak integration, and therefore higher confidence and precision of quantitative values eventually annotated to each peptide. We decided to focus on 50 SPD as the gradient length for all further experiments within this study as it shows the best proteome coverage with still 5 datapoints per peak on median (MS1 level), enough for proper integration. We further decided to quantify on the MS1 level as we not only see more datapoints there but also a lowered median Coefficient of Variation (CV) clearly below 10 %.

Next, we created a tailored library from 10 ng of the very same HeLa and K562 digests to further improve proteome coverage. The libraries were recorded in DIA mode as well and resulted in more than 62 000 precursors identified within the library (Figure 2). Using these libraries, we performed a library search for the 250 pg runs, which improved protein group identifications by 41.5 % and 31.7 % compared to a library-free search in method evaluation mode and by 26.3 % and 25.6 % compared to the library-free search with matching for HeLa and K562 respectively. We identified more than 7,500 proteins from a single run with as little as 250 pg of HeLa peptide input and more than 6,800 proteins with 250 pg of K562 peptides. These numbers increased further to a total of 7,800 and 7,060 unique protein groups identified from three replicates of HeLa and K562, respectively (Supplemental Figure 1 A, B). Of note, we identified a total number of 6,126 protein groups from HeLa using directDIA+ in method evaluation mode and only 6,017 using directDIA+ with matching replicates, indicating a more stringent false discovery rate (FDR) control and improved data quality when activating matching (which is active per default in Spectronaut 18). Of those 6,017 total proteins, 5,989 were found in all replicates, indicating an excellent data completeness (Supplemental Figure 1 A). Importantly, our results depict not only a single hit wonder but were successfully reproduced on a second Orbitrap Astral using a different batch for the analytical column and K562 digest (Supplemental Figure 1 C).

**Figure 2:**
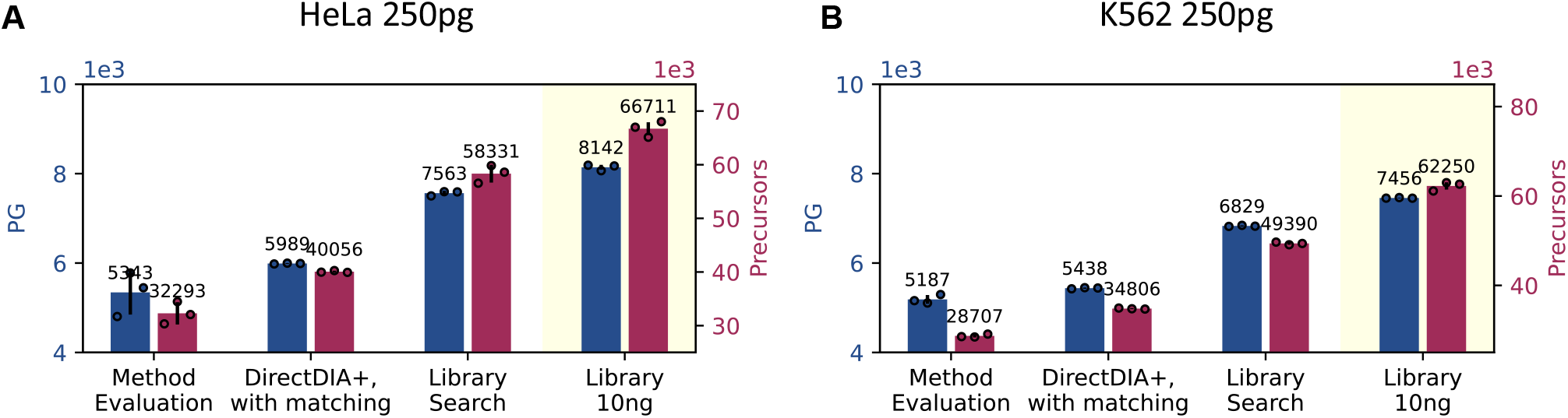
Influence of search strategy on the number of identified precursors and protein groups. 250 pg of HeLa (**A**) or K562 (**B**) peptides from diluted bulk digest were injected each. Peptides were separated at a throughput of 50 SPD. Data was recorded in DIA mode on the Orbitrap Astral MS and analyzed in the given mode using Spectronaut 18 with or without using a library from 10 ng peptides of the respective same cell-type, as indicated. Circles indicate identified protein groups (PG) or precursors at 1% FDR in individual replicates, bars indicate their means, while error bars indicate standard deviations with n=3. Of note, the last bar “Library 10ng” corresponds to the tailored library itself that we used for library search of 250pg runs.

### Matching across replicates does not negatively affect FDR

To estimate the real false discovery rate, we generated a shuffled target database by shuffling all sequences of our target database while maintaining the positions of all protease cleavage sites (P, K and R). Both databases were used as targets in the subsequent search. They have less than 0.01 % shared peptides and a similar size, allowing for a fair FDR estimation, with very low risk of finding peptides in the shuffled target database that represent true positive hits (Figure 3 A). Of note, randomization of peptide sequences led to a slightly increased number of unique peptides in the shuffled target vs target database, which we opted for deliberately to ensure a conservative estimate of the FDR.

**Figure 3:**
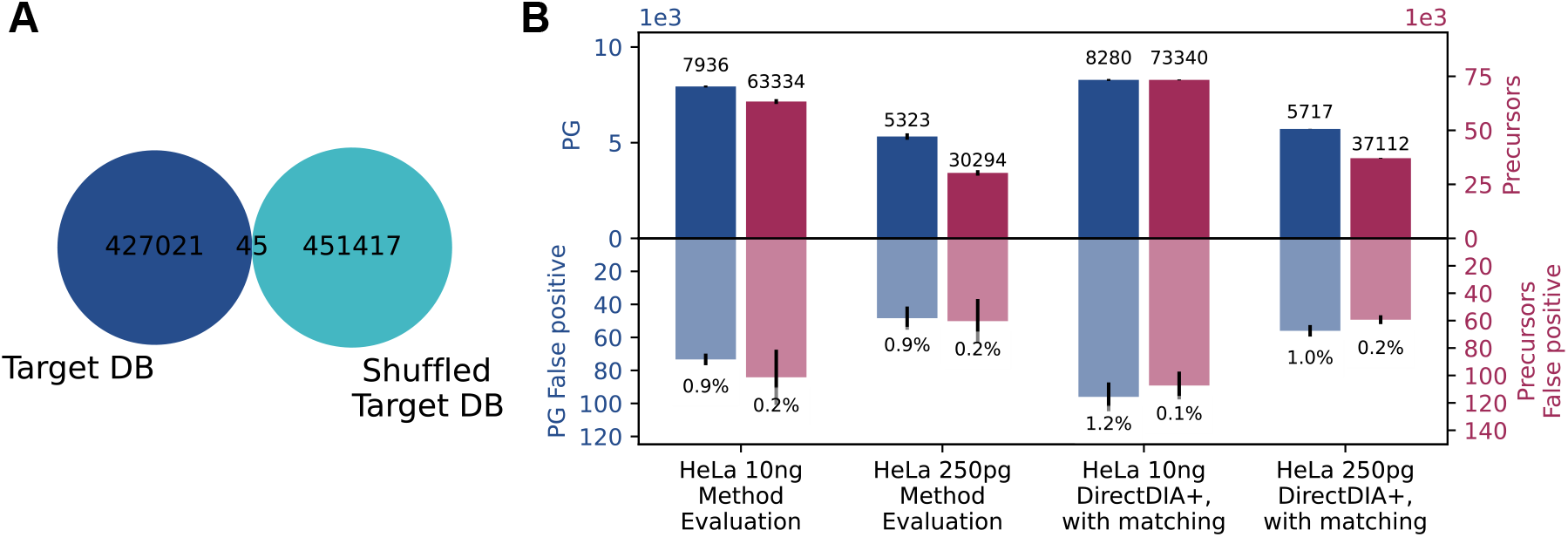
FDR dependence on search strategy and sample input amount. 250 pg or 10 ng of HeLa peptides from diluted bulk digest were injected each, reflecting the results shown in Figure 2, analyzed in the given mode using Spectronaut 18 at 1% FDR against a target (i.e., human proteome) and shuffled target (i.e., shuffled human proteome) database. **A**: Peptides of 8 – 30 amino acids length and assuming Leu = Ile contained in the target and shuffled target database. **B**: Bars indicate means of identified protein groups or precursors in the target database (top) and shuffled target database (bottom), while error bars indicate standard deviations with n=3.

When examining the HeLa data presented in Figure 2, we found that the FDR at protein group level is indeed below or very close to the expected 1 %, indicating an excellent FDR control using default settings in Spectronaut version 18 (Figure 3 B). This is in contrast to earlier results demonstrated in Spectronaut version 16 and 17, where FDR was reported to be higher than expected when using default parameters.^21^ When allowing for matching across replicates, the FDR only slightly increases to still acceptable levels with a maximum of 1.2 % false protein groups. On the precursor level, the FDR is very low and around 0.2 % for all conditions.

Reporting total ID numbers by summing up all unique IDs from 3 replicates (Supplemental Figure 1) without an additional FDR filter step, however, accumulates wrongly identified precursors and proteins. This leads to an elevated FDR of up to 2.7 % (Supplemental Figure 2). Hence, the authors advise reporting average ID numbers rather than total ID numbers.

### The FAIMS Pro interface improves S/N, CV and coverage

Aiming for the best possible sensitivity, we screened for the optimal compensation voltage using the FAIMS Pro interface. From previous experience, -48 V results in optimal results on the used device which is why that voltage was used for all other runs. Screening the compensation voltage from –38 to –88 V in steps of 10 suggests that the optimal setting is between -48 and -68 V, with few more protein groups and clearly more precursors detected at - 58 V compared to any other setting. While the FAIMS Pro interface usage bears the risk of losing some ions, it seems still advantageous to use it due to an improved signal to noise ratio and improved dynamic range.^22,23^ As already reported earlier by our lab^24^, a significant gain in sensitivity, especially for limited sample LC-MS/MS analyses, can be seen by using the FAIMS Pro interface, as the relative contribution of singly charged background ions gets more substantial at low sample loads.

In our hands, the FAIMS Pro interface usage at optimal settings resulted in a clearly improved signal to noise (Supplemental Figure 9), up to 42.6 % more precursors and 55.3 % more protein groups identified compared to measurements without a FAIMS interface (Figure 4 A). In line with our expectation from the improved signal to noise, we quantified more low abundant proteins when using FAIMS (Figure 4 D). However, some of the abundant precursors are lost, resulting in only 49% and 63% of peptides identified in runs without FAIMS can be found in the most promising compensation voltages -48V and -58V, respectively (Figure 4 E). And almost all (98%) peptides found in runs without FAIMS are present in one of the compensation voltages runs (Supplemental Figure 10). The observed effect is less dramatic on protein level, where close to all proteins quantified without FAIMS were also quantified with FAIMS. In addition, FAIMS usage results in more identified proteins. Global sequence coverage with FAIMS is slightly reduced (i.e., on average 7.8 peptides/ protein were identified without FAIMS but only 7.6 and 6.0 peptides/protein at a compensation voltage of -48 and -58 respectively). The acquisition speed was also not affected as there was no influence on the number of datapoints per peak (Figure 4 C) and the coefficient of variation (Figure 4 B) was strongly improved by FAIMS. Overall, the authors believe that FAIMS usage is highly advantageous for most single-cell studies mainly due to the improvement of signal to noise and CV that results in more proteins being identifiable- and quantifiable.

**Figure 4:**
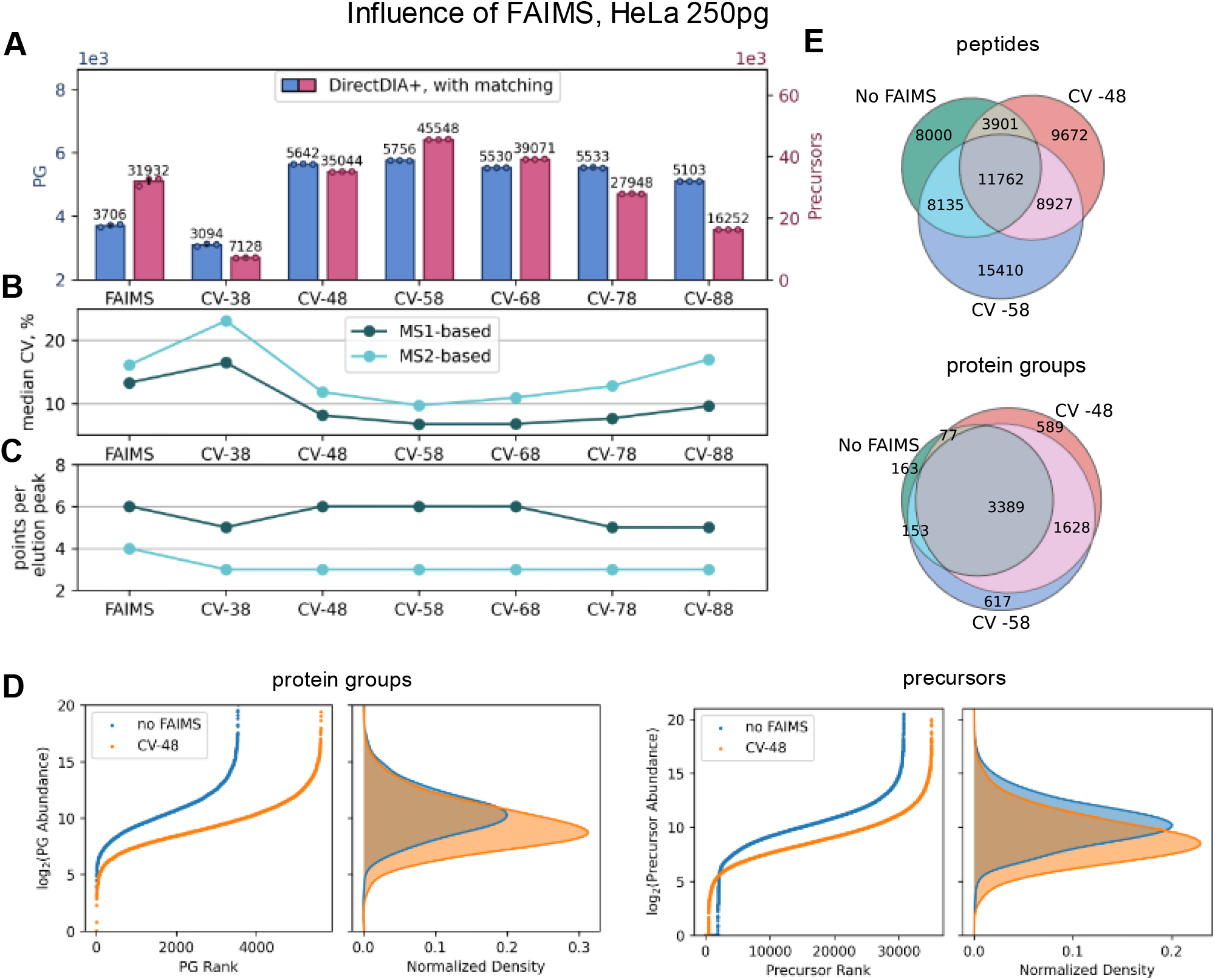
Influence of FAIMS compensation voltage on number of IDs and quantitative precision. 250 pg of HeLa peptides from diluted bulk digest were injected each. Peptides were separated at a throughput of 50 SPD. Data was recorded with or without the FAIMS Pro interface unit attached and at the given compensation voltage. **A**: Circles indicate identified protein groups (PG) or precursors at 1% FDR in individual replicates, bars indicate their means, while error bars indicate standard deviations. **B**: Dots indicate the median coefficient of variation on protein level when performing quantification on MS1 or MS2 level using Spectronaut 18. **C**: Dots indicate the median number of datapoints recorded per precursor when performing on MS1 or MS2 level as indicated. **D**: Log_2_ abundance ranks and their density plots for protein groups (left) and precursors (right) **E**: Venn diagrams show the number of protein groups or peptides quantified in 1 representative replicate each using either no FAIMS or FAIMS with a compensation voltage of -48 or -58. **A-C**: n=3.

### The Orbitrap Astral MS’ speed and sensitivity yield superior quantitative precision and accuracy over the Orbitrap Exploris 480 MS

We next evaluated the quantitative performance of our workflow at single-cell level inputs and benchmarked the data to our Thermo Scientific™ Orbitrap Exploris™ 480 MS using similar settings, the same input, and the same gradient length.

While the coefficient of variation gives a reasonable estimate of the quantitative precision across technical replicates, we further evaluated the accuracy and precision of proteins quantified from a two-proteome mix (Figure 5). By comparing two proteome mixes with different, but known, human to yeast mixing ratios we assessed the accuracy of quantification. Of note, less than 50 pg yeast, present in one of the samples, still enabled the quantification of 1782 yeast proteins. As shown in Figure 5 A, the Orbitrap Astral MS does an excellent job delivering a fold change in protein abundance very close to the expected one and outperforms the Orbitrap Exploris 480 MS (Figure 5 C), especially for the more sparse yeast samples. We further checked on accuracy by looking at the distribution around the expected fold change and found that higher abundant proteins within our sample tend to be quantified very accurately with a local CV of less than 5 %. The local CV, however, rises with lowered protein abundance and approaches 35 % at the limit of quantification. We observe an exponential drop of CV with increasing protein abundance (top panels, Figure 5).

**Figure 5:**
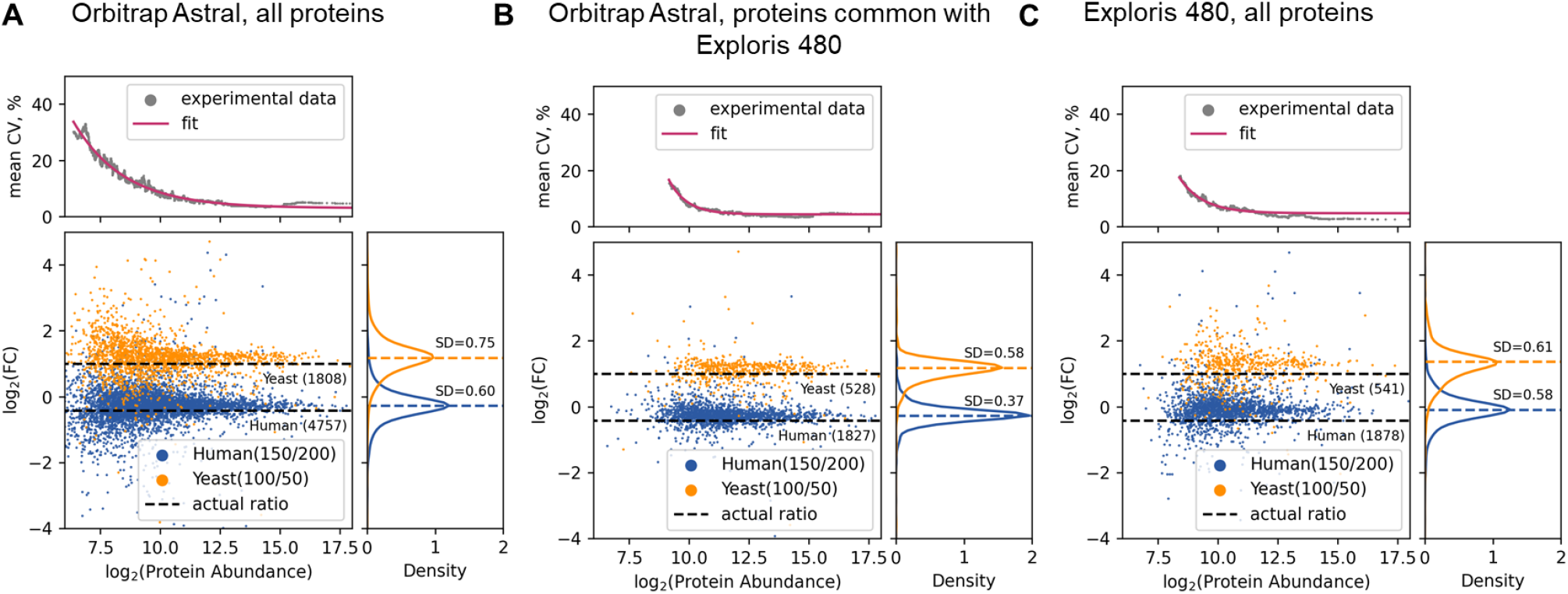
Human – Yeast proteome mix to assess quantitative precision and accuracy. From diluted bulk digests 250 pg two-proteome mixes consisting of either 150 pg HeLa + 100 pg yeast or 200 pg HeLa + 50 pg yeast were injected each. Peptides were separated at a throughput of 50 SPD. Data was recorded in DIA mode using optimal but not same settings for the Orbitrap Astral MS and Orbitrap Exploris 480 MS (see methods section) and analyzed using direct DIA+ in Spectronaut 18 at 1% FDR. Quantification was done on MS1 level. Dots within the Bland–Altman plots (bottom) represent proteins with given log2 average abundance and log2 fold change of abundance across both proteome mixes. Density plots (right)depict distribution of measured log2 fold changes and CV diagrams (top) show the local CV of 100 proteins quantified with a rolling window over the entire abundance range, n=3. **A**: Data obtained from the Orbitrap Astral MS showing all quantified proteins or, **B**: only those proteins that were commonly quantified in using the Orbitrap Exploris 480 MS. **C**: Data origiating from the Orbitrap Exploris 480 MS, showing all quantified proteins.

Our results show that on the Orbitrap Astral MS more than twice the number of proteins are quantified compared to the Orbitrap Exploris 480 MS, which is likely due to its improved speed and sensitivity. When filtering for proteins commonly found (Figure 5 B), predominantly lower abundance proteins are missing, highlighting the increased sensitivity of the Orbitrap Astral MS. Furthermore, almost all proteins identified with the Orbitrap Exploris 480 MS were also identified using the Orbitrap Astral MS. The number of common proteins is very close to the protein number quantified using the Orbitrap Exploris 480 MS. The commonly quantified proteins allow for a fair comparison of accuracy & precision between the Orbitrap Astral MS and the Orbitrap Exploris 480 MS and shows that the distribution of fold changes is clearly smaller on the Orbitrap Astral MS. The same is true for the local CV, most distinct for the lowest abundant proteins, where the CV drops from ∼20 % for the Orbitrap Exploris 480 MS to ∼12% for the Orbitrap Astral MS. Since quantification was done on MS1 level, data from both instruments originates from an Orbitrap analyzer of the same construction type. We hypothesize that the increased CV even for the very same proteins is due to the lower speed of the Orbitrap Exploris 480. The Orbitrap Exploris MS uses the Orbitrap analyzer for both the MS1 and MS2 data acquisition, whereas the Orbitrap Astral MS simultaneously utilizes the Orbitrap analyzer for MS1 and the Astral analyzer for MS2, which gives the latter one an enormous advantage in speed. This is also reflected in more data points per elution peak recorded on the Obritrap Astral MS (median 3 for Exploris and 5 on the Astral for this dataset).

Applying an additional filter to exclude proteins quantified based on only one peptide has no impact on the accuracy, but further improves the quantitative precision, independent of the instrument used (Supplemental Figure 4). As a tradeoff, 17 % (Orbitrap Astral MS) – 28 % (Orbitrap Exploris 480 MS) less proteins are quantified. We investigated both, accuracy and precision using larger fold changes of 5:1 and 10:1 in yeast quantity as well (Supplementary Figure 5), showing similar trends as aforementioned.

We believe that this data gives valuable input also for future biological applications aiming to estimate the minimal fold change regulation that is needed to be confidently identified by means of single-cell proteomics.

### Proof of principle study using single cells

#### Benchmarking label-free single-cell sample preparation workflows

To decide on an optimal label-free sample preparation strategy using the cellenONE robot, we benchmarked some protocols established on that machine. We decided to use the one-pot 384 well sample preparation published earlier in our group^7^ as a reference point and compared it to the proteoCHIP LF 48^25^ as well as to the recent proteoCHIP EVO96 protocol of Cellenion.^26^ While the one-pot 384 well workflow has a working volume of 1 μL this can be reduced to 300 nL in the proteoCHIP workflows enabling a higher sample and protease concentration that we expect to reduce losses and reduce excess trypsin needed. As a tradeoff, a direct injection from the proteoCHIP to the LC-MS system is tricky and bears the risk injecting hexadecane into the MS system (in case of the LF 48 workflow, no hexadecane was used in the EVO96 chip). This requires a sample transfer step that is done either by manual pipetting (LF48) or by centrifugation (EVO96) to a 96 well low binding PCR plate.

In our first attempt, the proteoCHIP LF48 by far outperformed the one-pot 384 well workflow by means of 48 % more protein groups identified. However, out of 1979 protein groups identified using the LF48 chip, 693 (35 %) were identified in a blank control, that contained all reagents and received the same treatment as single cell samples, but no cell was added (Supplemental Figure 3 A). To reduce the background, we implemented a stringent washing protocol for the Teflon LF48 chip. As a result, the background level of our blank control was reduced to the same level as seen for 384 well plates but the ID numbers were reduced to 1128 protein groups, slightly less than what we obtained using the 384 well plate (1341 protein groups). We hypothesize that contamination peptides covered the (non-stringent cleaned) surface of the (multiple used) LF48 chips serving as a carrier, but potentially covering real (regulated) proteins.

Next, we compared the more recent (multiple used and washed) EVO96 chip prototype made of polypropylene, to the one-pot 384 well protocol, yielding more identified protein groups even at lower background for the 384 well plate. Selecting only big (>20 μm) A549 cells yielded 3690 protein groups on average using the one-pot 384 well workflow, compared to 2348 protein groups using the EVO96 chip with a single sample transfer to a 96 well low binding plate (Supplemental Figure 3 B).

We conclude that, in our hands, the one-pot 384 workflow seems best suited for sample preparation due to competitive or even superior ID numbers and simpler handling as no transfer step or removing of hexadecane is needed. Furthermore, reporting blanks seems of high importance to us to check on background contamination, that is especially problematic in case of multiple usage of chip materials. All single cell samples in this study were therefore prepared using the one-pot 384 well^7^ workflow.

#### Study of A549 cells

A549 is a human non-small cell lung cancer cell line commonly used for basic research and drug discovery. We decided to use this cell-line for proof of principle studies with our workflow. Three different size groups ranging from 15-30 μm diameter were collected using the cellenONE robot. 66 single A549 cells and 3 blanks were measured using the Orbitrap Astral MS. As expected, the number of protein groups identified correlates with the size of the cells (Figure 6 A) and ranges from 1801 protein groups from small cells to 2870 protein groups for large cells on average when searching library free and in method evaluation mode. When allowing for matching across the single cells, ID numbers are improved to 3304 (+83%) and 4439 (+55%) for smaller and bigger cells respectively. We investigated if the use of a spectral library might further increase ID numbers and created a tailored library either from 20x or 40x A549 cells isolated into a single well of the 384 well plate, processed and measured using the exact same settings as for single cells. We could not significantly improve proteome coverage using a library search instead of directDIA+, although the used 20x and 40x libraries contained much more proteins than found in single cells (6395 protein groups for 20x and 6762 protein groups for 40x). Using the 40x library, ID numbers even dropped a bit (Figure 6 A). This is why we assume there is a sweet spot in optimal library size: MS acquisition parameters, optimized for a single cell input, as accumulation time or m/z windows, are likely not perfectly optimized anymore for bigger amounts as used for a large library. In addition, we believe that the high number of replicates already serves as an excellent basis for matching. In line with those assumptions, we observed a more pronounced effect of using a library when optimizing the MS method for the higher input and when using less replicates of single cells, as done in our second study on TE-like cells and naïve hPSC cells (Figure 7). In our hands, using a 20x library yielded the best results and enabled us to identify up to 5300 proteins from a single A549 cell.

**Figure 6:**
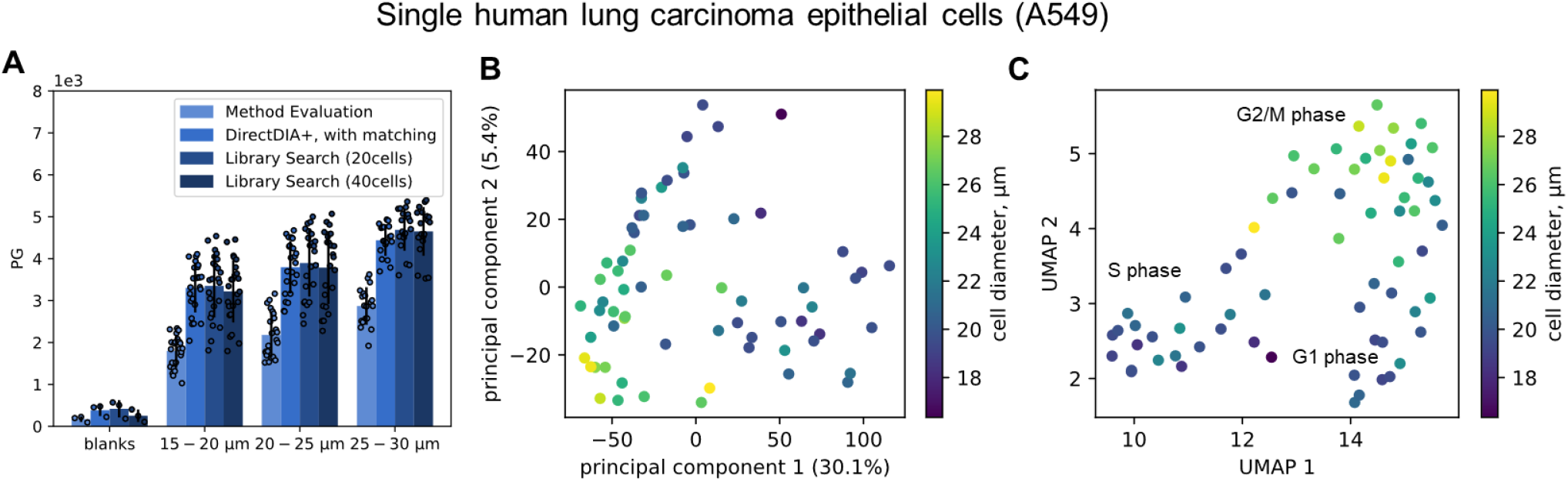
Analysis of single A549 cells of different sizes. Individual A549 cells were prepared^7^ in a 384 well plate using the cellenONE in predefined size groups of 15-20, 20-25 or 25-30 μm diameter. Digested cells were analyzed at 50 SPD using the previously optimized settings for LC and MS on the Orbitrap Astral MS. **A**: Circles indicate identified protein groups (PG) at 1% FDR of each individual cell, bars indicate their means, while error bars indicate standard deviations. The search was performed library free in method evaluation mode, or with matching, or against a tailored library created from 3 replicates of 20x or 40x cell, all samples were prepared and measured in the exact same way. Blanks were processed in the same 384 well plate and contained all reagents and buffers but no cell. **B**: PCA and **C**: UMAP based on protein quantities with each dot representing a cell and colors reflecting the actual cell size as determined by the camera system of the cellenONE.

**Figure 7:**
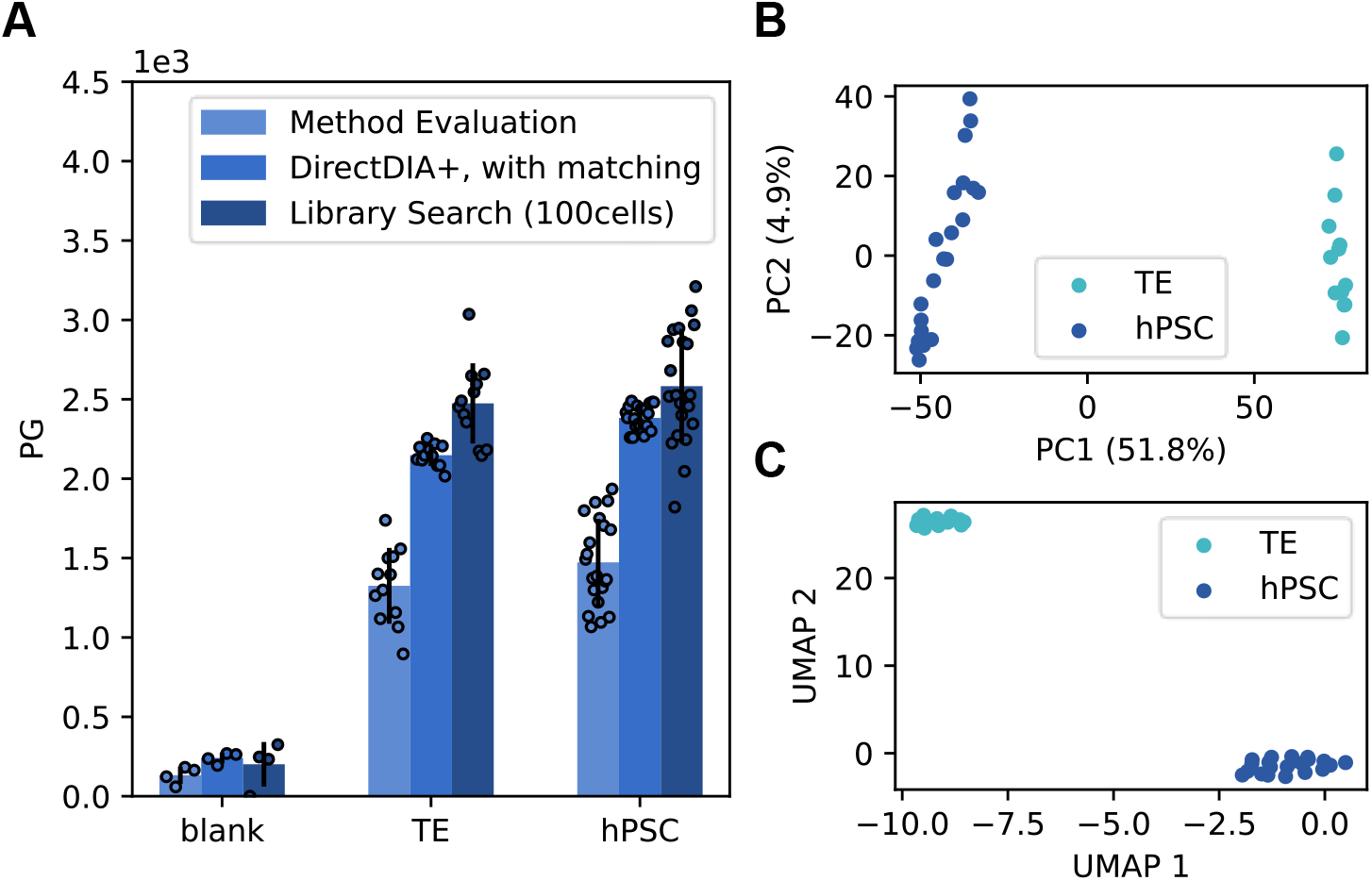
Analysis of single TE and hPSC cells. Individual cells were isolated using FACS into a 384 well plate. Digested cells were analyzed at 50 SPD using the previously optimized settings for LC and MS on the Orbitrap Astral MS. **A**: Circles indicate identified protein groups (PG) at 1% FDR of each individual cell, bars indicate their means, while error bars indicate standard deviations. Search was performed library free in method evaluation mode or with matching or against a tailored library created from 3 replicates of 100x cells prepared in the exact same way but recorded with adopted settings (i.e., 60 ms I.T. and m/z 20 windows for single cells, 10 ms I.T. and m/z 5 windows for 10 cells. Blanks were processed in the same 384 well plate and contain all reagents and buffers but no cell. PCA (**B**) UMAP (**C**) analysis based on protein quantities with each dot representing a cell and colors reflecting the cell type based on their fluorescent marker proteins.

We next examined the extent of cellular heterogeneity of these cultured and untreated A549 cells and found cells clustering differently dependent on their diameter by principal component analysis (PCA, Figure 6 B). This is even more pronounced when doing a uniform manifold approximation and projection (UMAP, Figure 6 C). We took a deeper look at the most abundant proteins identified in our single cells, which were the histone H4 (H4C1) and the actines ACTB and ACTC1. Indeed, their abundance profiles are nicely reflected in the clustering seen in the PCA or UMAP (Supplementary Figure 6). We suspect that their abundance differences might not only correlate to the recorded cell sizes, but that both might correlate to different cell cycle stages. G2/M phase cells were reported to be of biggest size^27^, which is why we annotated our biggest cell size cluster as presumably G2/M. Next, we tried to confirm this by checking the expression pattern of abundant and known marker proteins. Actines are known to play a crucial role in cell division, for cell junctions and cell shape, highly relevant especially for cancer cells, which is why it seems logical that their expression levels correlate to the cell size and cell cycle stage.^28,29^ Histone levels are known to be elevated during G2/M phase when DNA synthesis takes place. To check if those cells with an elevated H4C1 and H14 level in our PCA and UMAP plots (Supplementary Figure 6) could indeed be primarily in G2/M phase we investigated the abundance levels of the nuclear ubiquitous casein and cyclin-dependent kinase substrate 1 (NUCKS1) that was also quantified in our dataset. NUCKS is known to be expressed at high levels in S phase^30,31^ and is indeed present at lowered expression levels in those cells presumably in G2/M phase but upregulated in another cell cluster (presumably in S phase). We additionally checked for a classical cell cycle marker (CDK1). CDK1 was reported to have aligned expression levels to those of NUKS1^30^, but we could not see a clear trend in our data. MKI67, another reporter is expected to be maximally expressed in G_2_ phase^32^, also aligning with or results and correlating to the size distribution (Supplemental Figure 6, Figure 6 B).

Of note, no cell cycle control was performed in this study, which is why our cellular population reflects a wild mix of all phases, impeding a clear separation but still allowing to successful study the heterogeneity of this system.

### Characterization of human hPSC and TE-like derivative

To challenge our analytical LC-MS/MS setup and the abilities of SCP further we analyzed naïve hPSCs and induced from them TE-like cells. On day 4 TE-like cells were harvested and sorted for TROP2+ using FACS. We evaluated different analysis strategies (Figure 8 A). With our library-based approach, we could identify 2339 and 2544 protein groups in TE-like cells and naïve hPSCs, respectively.

To confirm that SCP can recapitulate known differences between naïve hPSCs and TE-like cells, we performed dimensionality reduction on all cells in the dataset using linear PCA and non-linear UMAP clustering analysis. We observed strong separation between these two cell populations (Figure 8 B, C) with both approaches, which aligned with cell types. PCA reported an explained variance of 51.8% (PC1) and 4.9% (PC2).

Gene ontology (GO) analysis of proteins upregulated in TE-like cells showed enrichment in the following biological processes: cytoskeleton organization (GO: 0007010), actin cytoskeleton organization (GO: 0030036),, epithelial cell differentiation (GO: 0030855), epithelium development (GO:0060429), positive regulation of cell differentiation (GO:0045597), cell differentiation (GO:0030154), which rightfully corresponds to the previously described biological processes underlying the differentiation of hPSC to TE-like cells^33^ (Supplementary Figure 8).

TE cell analogues were identified as *GATA2, GATA3, CDX2*, and *TROP2* positive.^34^ Transcription factors are typically low abundant in the cell and are often challenging to detect with proteomics techniques. They are also partly localized in the nucleus, which can make them less accessible for extraction and analysis. Additionally, many transcription factors are modified by phosphorylation or other post-translational modifications that can affect their detection. Here, we were able to identify GATA2 and GATA3 at very low abundance in some TE-like cells. Among differentially expressed proteins between hPSCs and TE-like cells, we found many TE markers, well known from transcriptome data: KRT18 (epithelium cytoskeleton), KRT19 (epithelium cytoskeleton), RAB25 (RAS pathway), DAB2 (ERK pathway), YAP1 (Hippo pathway), S100A16, SP6, CDH1, ENPEP, SLC7A2, HAVCR1, PDLIM1 — associated with TE development; and DPPA4, DPPA2, SUSD2, DNMT3L which are well established naïve hPSC markers^33,35^ (Supplementary Figure 7). Abundances for these proteins correlate with transcriptome data known from literature.^33,35^

Our single-cell proteome data of naïve hPSCs and TE-like cells demonstrates the capacity to distinguish primary cells and to extract meaningful biological information that complements and enriches results from transcriptome data.

## DISCUSSION

Despite tremendous improvements already made in the field of ultra-low input proteomics by mass spectrometry in the past decade, achieving a high sample throughput, reproducibility, quantitative accuracy, precision, and high sensitivity at the same time is still very challenging. The presented workflow aims to fulfill those criteria by combining a high performance zero dead end volume column at a short gradient with one of the most advanced, fast, and sensitive MS instruments, FAIMS based noise reduction and a DIA-based acquisition. The used packed bed column has a pulled emitter directly at the end of the column, which minimizes peak broadening, and which is further supported by the usage of high flowrates during sample loading, increasing the throughput. LC flowrates are reduced during the active gradient to enhance sensitivity. We found that total throughputs of 50 SPD represent a sweet spot between maximizing speed of sample acquisition and maintaining enough measurement time to detect a median of 5 data-points per elution peak enabling proper quantification. We further found that there is another sweet spot of optimal compensation voltage when using the FAIMS Pro interface at -48 to -58 V. Noise was reduced and sequence coverage for those proteins also identified without FAIMS was not negatively affected. Total proteome coverage was at the same time enhanced, which is why using FAIMS for limited amount samples is clearly advantageous in our hands.

We further found that a tailored library and adopted isolation window sizes are advantageous to maximize precursor and protein identification numbers. This yielded in surprisingly high numbers close to 7,400 protein groups identified from as little at 250 pg HeLa and up to 5,200 from a single A549 cell. FDR was not negatively affected by allowing matching and could be confirmed to be very close to the expected 1 % using default settings in Spectronaut 18.

To the best of the author’s knowledge, we are the first to assess quantitative accuracy and precision at single cell level using a two-proteome mix. We found that quantitation works surprisingly well and that 5 data points per elution peak as reached on the Orbitrap Astral MS at 50 SPD seem sufficient for successful separation of 2-fold changes in protein abundance. However, the CV of fold changes from quantified proteins within a sample increase with decreased protein abundance and ranges from quite low 5 % up to 40 % for the lowest abundant proteins within our proteome mixes. Hence, relative abundance within a single cell sample is an important consideration also for future biological studies as the confident clustering of cellular populations currently can only rely on more abundant proteins within single cells. Despite this limitation, the Orbitrap Astral MS clearly outperforms the Orbitrap Exploris 480 MS in terms of number of quantified proteins, sensitivity, and quantitative accuracy, allowing to investigate heterogeneity of untreated cultured cells based on their size and cell cycle phase, as shown for A549 cells. Furthermore, the distinct separation we observed between TE-like cells and naïve hPSC cells highlights the potential of this SCP workflow for application to human blastocysts. It could significantly aid in exploring TE development mechanisms, blastocyst morphology regulation, and the identification of molecular markers of high-quality blastocysts. Ultimately, this knowledge could contribute not only to our understanding of human blastocyst development but also potentially improve IVF success rates.

In conclusion, we are convinced that our comprehensive workflow with optimized and improved parameters from sample preparation to data interpretation is a highly valuable contribution to evolve single-cell proteomics to the next level in terms of sensitivity, reproducibility, and quantitative performance helping to transition the field of single-cell proteomics, from developmental phase to a technique for the biologist’s toolbox.

## METHODS

### Cultivation of A459 and HeLa cells

Cells were cultured at 37 °C in a humidified atmosphere at 5% CO_2_. A549 cells were grown in Roswell Park Memorial Institute (RPMI) 1640 Medium and HeLa cells grown in Dulbecco’s Modified Eagle’s Medium (DMEM). Cell medium was supplemented with 10% FBS (10270, Fisher Scientific, USA), 1x penicillin-streptomycin (P0781-100ML, Sigma Aldrich, Israel), 100X L-Glut 200 mM (250030-024, Thermo Scientific, Germany) and 1 mM Sodium-pyruvate (for RPMI only; 4275, Sigma-Aldrich, UK). Cells were grown to around 75% confluency before trypsinization with 0.05% Trypsin-EDTA (25300-054, Thermo Scientific, USA), followed by washing 3x with phosphate-buffered-saline (PBS). Cells were resuspended in PBS at a density of 200 cells/μL for isolation with the cellenONE®.

### Single cell isolation and sample preparation of A459 and HeLa cells

A549 and HeLa cell isolation, lysis and digestion was performed within a 384-well plate (Thermo Scientific™ Armadillo 45PCR Plate, 384-well, # 12657516) using the cellenONE® robot as previously described.^7^ Briefly, cells were sorted into wells containing 1μL of master mix (0.2% n-dodecyl-beta-maltoside (D4641-500MG, Sigma Aldrich, Germany), 100mM tetraethylammonium bicarbonate (17902-500ML, Fluka Analytical, Switzerland), 3ng/μL trypsin (Trypsin Gold, V5280, Promega, USA) and 0.01% enhancer (ProteaseMAX, V2071, Promega, USA) and 1 % DMSO). For single cell samples, cells were deposited into individual wells, while for 20x or 40x libraries, the respective cell-number was sorted into a single well. Humidity and temperature were controlled at 50% and 15°C during cell sorting. A549 cells were isolated at given diameter of 15-30 μm and HeLa cells at 18-25 μm. The maximum elongation was set to 1.5. Cell lysis and protein digestion were done at 50°C and 85% relative humidity for 30 min before an additional 500nL of 3ng/μL trypsin were added. After lysis and digestion, 3.5 μL of 0.1% TFA were added to the respective wells for quenching and storage at -20°C. For the LC-MS/MS analysis, samples were directly injected from the 384 well plate.

For benchmarking studies to the proteoCHIP® based sample preparation, cells were isolated and prepared exactly as described in the manufacturer’s manual. Standard (non-stringent) washing of chips was performed by sonication in methanol for 20 minutes followed by extensive flushing with milliQ water and drying in the fume hood. A stringent washing protocol was established by Cellenion and is considered a company secret. In brief, 300 nL of mastermix (0.2% DDM, 10 ng/μL Trypsin Gold, 100 mM TEAB) were dispensed into each well of the LF48 proteoCHIP® or the EVO96 proteoCHIP® (Cellenion, Lyon, France) using the cellenONE® robot. The LF 48 chip was prefilled with 2 μL hexadecane oil (H6703-100ML, Sigma-Aldrich, Darmstadt, Germany) per well. ProteoCHIPs were cooled to 8°C for cell isolation which freezes the hexadecane (LF 48). Cells were isolated as described above, following stepwise heating to final 50°C at 85% relative humidity for lysis and digestion. Constant addition of water by the cellenONE® keeps the samples hydrated and at the approximate constant volume. After that, 3.5 μL 0.1% TFA was added and the proteoCHIP LF 48 was cooled on wet ice to freeze the hexadecane again and allowed for manual separation from the sample containing aqueous phase by transferring the sample to individual wells of a low binding 96 well PCR plate (EP0030129512, twin.tec. PCR Plate 96 LoBind, skirted, Eppendorf, Darmstadt). The EVO96 proteoCHIP did not contain hexadecane and was directly placed on top of a 96well PCR plate to transfer the sample by centrifugation (500 g, 30 sec).

### Naive hPSC cell culturing

Naive PSCs were cultured on gelatin-coated plates with a feeder layer of gamma-irradiated mouse embryonic fibroblasts (MEFs). The coating and feeder layer preparation details were as per previously reported methods.^33^ Cells were cultured in PXGL medium, composed of N2B27 basal medium supplemented with 1 μM PD0325901 (Med-ChemExpress, HY-10254), 1 μM XAV-939 (MedChemExpress, HY-15147), 2 μM Gö 6983 (MedChemExpress, HY-13689), and 10 ng/ml human leukemia inhibitory factor (hLIF, in-house made). The N2B27 basal medium formulation included 50% DMEM/F12, 50% neurobasal medium, N-2 and B-27 supplements, GultaMAX supplement, non-essential amino acids, and 100 μM 2-mercaptoethanol. Cells were maintained in a hypoxic chamber (5% CO2, 5% O2), and passaged every three to four days. All cell lines were routinely tested negative for mycoplasma.

### TE differentiation from naïve hPSC

Naive PSCs were dissociated using Accutase (Biozym, B423201) at 37°C for 5 minutes. Gentle mechanical dissociation was performed using a pipette, followed by centrifugation to collect the cell pellet. The pellet was resuspended in PXGL medium, supplemented with 10 μM Y-27632 (MedChemExpress, HY-10583). To exclude MEFs, the cell suspension was transferred onto gelatin-coated plates and incubated at 37°C for 70 minutes. Cell counting and viability were determined using a Countess automated cell counter (Thermo Fisher Scientific) with trypan blue staining. Cells were seeded at a density of 1.2×105 cells per well on Geltrex (0.5 μl/cm2)-coated 6-well plates. Culturing was performed in a hypoxic chamber (5% CO2, 5% O2). On induction day 0, the medium was replaced with PDA83 medium comprising N2B27 basal medium, 1 μM PD0325901, 1 μM A83-01 (MedChemExpress, HY-10432), and 2% FBS. On day 1, 2 μM XMU-MP-1 was added. Days 2 and 3, the medium was switched to N2B27 with 2% FBS. On day 4, FACS sorting was conducted using the following protocol.

### TE and hPSC FACS sorting

TE-like cells and naïve PSCs were dissociated using Accutase at 37°C for 10 minutes and 5 minutes, respectively. Gentle mechanical dissociation was performed using a pipette. Cells were then stained with antibodies against TROP2 and SUSD2, respectively. For sorting TE-like cells, TROP2+ cells were selected and sorted into 384-well plates, ensuring exclusion of non-differentiated cells. Similarly, for naïve hPSC sorting, SUSD2+ cells were sorted into 384-well plates to exclude MEF. In both conditions, dead cells were identified and excluded using DAPI staining.

### Two-proteome mixes

HeLa (H) (Thermo Scientific, Pierce™ HeLa Protein Digest Standard, 88328) and yeast (Y) (Promega, MS Compatible Yeast Protein Extract, Digest, Saccharomyces cerevisiae, 100ug, V7461) were combined in 0.1% TFA at the following ratios and at a concentration of 250 pg/μL: H:Y = 200 pg : 50 pg; 240 pg : 10 pg; 150 pg : 100 pg. 1 μL (=250 pg) of each two-proteome mix were injected to assess limits of quantification on at ultra-low input levels.

### Diluted bulk digests

HeLa (Thermo Scientific, Pierce™ HeLa Protein Digest Standard, 88328) or K562 (Promega, Mass Spec-Compatible Human Protein Extract, V6951) were dissolved in 0.1% TFA, 0.015 % DDM (N-Dodecyl β-D-Maltosid, D4641-500MG, Sigma Aldrich, Germany) at a concentration of 5 ng/μL for injection into the LC-MS system.

### LC-MS analysis

Samples were analyzed using the Thermo Scientific™ Vanquish™ Neo UHPLC system (Thermo Scientific). Peptides were separated on an Aurora Ultimate TS 25 cm nanoflow UHPLC column with integrated emitter (Ion Optics, Fitzroy, Australia) at 50°C using direct injection mode.

Peptide separation was performed at 30 – 80 SPD with details given in Supplemental Table 1 and with all single cell measurements performed at 50 SPD. Fast sample loading was performed at a maximum pressure of 1400 bar and at a maximum flow of 1μL/min.

For MS measurement the Orbitrap Astral mass spectrometer or, for benchmarking on an Orbitrap Exploris 480 Mass Spectrometer (both Thermo Scientific) was coupled to the LC. Both instruments were equipped with a FAIMS Pro interface (Thermo Scientific) and an EASY-Spray source. Data recorded for reproducibility checks (Supplemental Figure 1) was recorded on a second Orbitrap Astral instrument equipped with a FAIMS Pro Duo interface. A compensation voltage of -48 (Orbitrap Astral) or -50 (Orbitrap Exploris 480) was used, if not indicated differently. An electrospray voltage of 1.85 kV was applied for ionization and adopted to slightly higher voltages for aged emitters to ensure spray stability.

On the Orbitrap Astral MS, MS1 spectra were recorded using the Orbitrap analyzer at 240k resolution from m/z 400-900 using a AGC target of 500% and a maximum injection time of 100 ms. For MS2 in DIA mode using the Astral analyzer, non-overlapping isolation windows ranging from m/z 5 (100 cells, 10 ng) to m/z 20 (1-40 cells, blanks, 250 pg bulks) and a scan range from m/z 400 – 800 were chosen. The precursor accumulation time ranged from 10 ms (100x cells, 10 ng bulks) to 40 ms (250 pg bulks) to 60 ms (1-40 cells, blanks) and the AGC target was set to 800%.

On the Orbitrap Exploris 480 MS the MS method was altered due to the lowered speed and sensitivity of that instrument compared to the Orbitrap Astral MS. However, comparable settings for high throughput analysis as tested earlier^12^ were used with the following details: MS1 spectra were recorded using the Orbitrap analyzer at 120k resolution from m/z 400-800 using a AGC target of 300%. For MS2 in DIA mode, overlapping (m/z 1) isolation windows of m/z 40 and a scan range from m/z 400 – 800 were chosen. The precursor accumulation time was set to a maximum of 118 ms and the AGC target was set to 1000%.

### Data analysis

All raw-data was analyzed using Spectronaut (v 18.6.231227.55695, Biognosys AG).^36^ For results marked as “Method Evaluation”, directDIA+ was used in method evaluation mode without cross normalization and with every raw file defined as separate condition to ensure Spectronaut treats each file like it they were analyzed individually. “Direct DIA+ with matching” indicates results analyzed in directDIA+ mode with replicates defined as same condition in the search settings. For library searches higher input DIA recorded results (as given) were first analyzed in direct DIA+ mode and a library was created from those files using Spectronaut.

Quantification was performed on MS1 level. Carbamidomethylation of cysteines as a static modification was removed for single-cell searches as no alkylation step was performed. Besides that, factory settings were used for all analyses and for library generation. Searches were performed against the human proteome (uniprot proteome UP000005640, reviewed, 20408 protein entries, downloaded 4^th^ Aug 2023) and the crapome^37^ (118 protein entries). Searches for the two-proteome analysis were run against the same human database and the yeast proteome (UniProt proteome UP0000023116050, reviewed, 6727 protein entries, downloaded 11^th^ Dec. 2023).

For FDR checks a decoy (“shuffled target”) database was generated using pyteomics^38^ Python package in shuffled mode with fixed positions for arginine, proline, and lysine to make sure that peptide mass and length distributions are the same as in a target database.

### Post Analysis

#### Human and Yeast dataset

Raw quantity values were used. All proteins with missing quantitative values were filtered out.

#### Single-cell datasets

All proteins with missing values in more than 20 cells for A459 and 5 cells for TE with hPSC dataset were filtered out. The rest of the missing values were substituted with intensity equal to minimum in the dataset. Then quantitative data were transformed to log2-format and normalized to normal distribution. The cell diameter data for PCA and UMAP clustering was taken from cellenONE robot. PCA and UMAP clustering were done using sklearn^39^ and umap^40^ Python packages.

To find statistical differences between hPSC and TE cells, a Student’s T-Test for two independent samples was performed. Benjamini-Hochberg FDR was employed to correct p-values for multiple comparisons. The level of significance for corrected p-values was set to 0.05. Proteins with fold-changes (TE/hPSC) more than one were considered as up-regulated. GO analysis of up-regulated proteins was performed using String-db.org.^41^

## Ethical approvals

The Wicell line H9 was used under the agreement 20-WO-341 for a research program entitled ‘Modeling early human development: Establishing a stem cell-based 3D in vitro model of human blastocyst (blastoids)’. Blastoid generation was approved by the Commission for Science Ethics of the Austrian Academy of Sciences. This work did not exceed a developmental stage normally associated with 14 consecutive days in culture after fertilization even though this is not forbidden by the ISSCR Guidelines as far as embryo models are concerned. All experiments complied with all relevant guidelines and regulations, including the 2021 ISSCR guidelines that forbid the transfer of human blastoids into any uterus.^42^

## ACKNOWLEDGMENTS

This work was supported by the infrastructure funding 4th call 2022/01 (AT-SCP) of the Austrian Research Promotion Agency (FFG) and the project LS20-079 of the Vienna Science and Technology Fund (WWTF). This work was further funded by the P35045-B project (Grant-DOI 10.55776/P35045) and the F 8801-B Meiosis project (Grant-DOI 10.55776/F88) of the Austrian Science Fund (FWF). This research was funded in whole, or in part, by the Austrian Science Fund (FWF). JAB’s work was funded by FWF (Grant-DOI 10.55776/ESP497). The authors thank the group of Josef Penninger at IMBA for providing A549 cells. For the purpose of open access, the authors have applied a CC BY public copyright license to any Author Accepted Manuscript version arising from this submission.

## AUTHOR CONTRIBUTIONS STATEMENT

JAB and MM conceptualized the study, designed, and performed experiments, data analysis, and wrote the manuscript. TNA, ED and BD performed experiments and designed experiment settings. JS and HK prepared TE and hPSC. JS, HK, TMS, and NR interpreted TE and hPSC data. PP contributed to the data analysis strategy. KM and MM supervised the study. All authors revised and agreed on the manuscript.

## DATA AVAILABILITY

The mass spectrometry proteomics data have been deposited to the ProteomeXchange Consortium via the PRIDE^131^ partner repository with the dataset identifier PXD049412. For review, data can be accessed via the following login: Username, Password: erFjvN8w

