## Supplemental Material for "Challenging the Astral^™^ mass analyzer - up to 5300 proteins per single-cell at unseen quantitative accuracy to study cellular heterogeneity"

* These authors contributed equally

### Supplemental Tables

**Supplemental Table 1:** LC gradient details, buffer B: acetonitrile with 0.08% formic acid, buffer A: 0.1% formic acid

| | 30 SPD | | 40 SPD | | 50 SPD | | 60 SPD | | 80 SPD | |
|---|---|---|---|---|---|---|---|---|---|---|
| time [min] | flow [nl/min] | % buffer B | flow [nl/min] | % buffer B | flow [nl/min] | % buffer B | flow [nl/min] | % buffer B | flow [nl/min] | % buffer B |
| **0** | 250 | 5 | 450 | 5 | 450 | 1 | 450 | 1 | 500 | 4 |
| **0.1** | - | - | - | - | 450 | 4 | 450 | 4 | 500 | 4 |
| **0.8** | 250 | 8 | 450 | 8 | - | - | - | - | - | - |
| **1.0** | - | - | 450 | 10 | - | - | - | - | - | - |
| **1.1** | - | - | 250 | 10.3 | - | - | - | - | - | - |
| **1.6** | - | - | - | - | - | - | - | - | 500 | 12 |
| **1.7** | - | - | - | - | - | - | - | - | 200 | - |
| **1.9** | - | - | - | - | 450 | 12 | 450 | 12 | - | - |
| **2** | - | - | - | - | 200 | - | 200 | - | - | - |
| **9.7** | - | - | - | - | - | - | - | - | 200 | 28.5 |
| **11.2** | - | - | - | - | - | - | - | - | 200 | 40 |
| **12** | - | - | - | - | 200 | 22.5 | - | - | 300 | 99 |
| **13.5** | - | - | - | - | - | - | 200 | 28.5 | - | - |
| **14** | - | - | - | - | - | - | - | - | 500 | 99 |
| **17** | - | - | - | - | - | - | 200 | 40 | - | - |
| **18** | - | - | - | - | - | - | 300 | 99 | - | - |
| **19.5** | - | - | - | - | 200 | 40 | - | - | - | - |
| **20** | - | - | - | - | - | - | 300 | 99 | - | - |
| **22** | - | - | - | - | 300 | 99 | - | - | - | - |
| **25** | - | - | - | - | 300 | 99 | - | - | - | - |
| **25.1** | - | - | 250 | 30 | - | - | - | - | - | - |
| **27.6** | - | - | 250 | 44 | - | - | - | - | - | - |
| **28.4** | - | - | 250 | 99 | - | - | - | - | - | - |
| **30.8** | 250 | 30 | - | - | - | - | - | - | - | - |
| **31.4** | - | - | 450 | 99 | - | - | - | - | - | - |
| **33.8** | 250 | 44 | - | - | - | - | - | - | - | - |
| **34.8** | 250 | 99 | - | - | - | - | - | - | - | - |
| **38.8** | 250 | 99 | - | - | - | - | - | - | - | - |

### Supplemental Figures

**Supplemental Figure 1:** Influence of search strategy on the total number of identified precursors and protein groups, and long-term reproducibility.

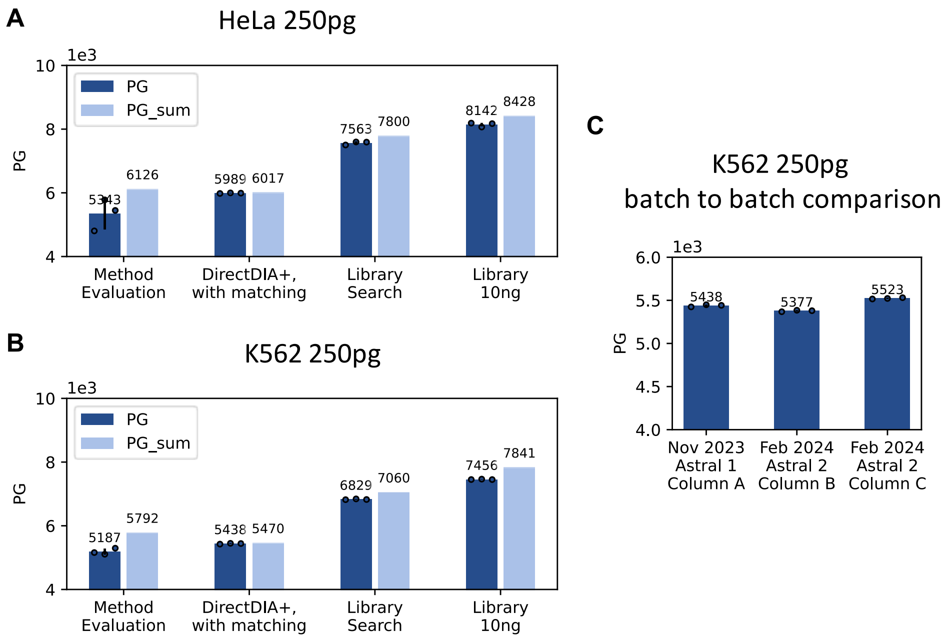

*__Supplemental Figure 1__: 250 pg of HeLa (A) or K562 (B,C) peptides from diluted bulk digest were injected each. Peptides were separated at a throughput of 50 SPD. Data was recorded in DIA mode on the Orbitrap Astral MS and analyzed using Spectronaut 18 with or without using a library from 10 ng peptides of the respective same cell-type, as indicated. __C__: Data was analyzed in directDIA+ with matching, comparing measurements from 2 batches (November 23 and February 24) on two different Orbitrap Astral MS and using 3 different columns. Circles indicate identified protein groups (PG) or precursors at 1% FDR in individual replicates, bars indicate their means (dark blue) or the total number of unique identifications in 3 replicates (light blue), while error bars indicate standard deviations with n=3.*

**Supplemental Figure 2:** FDR on total number of identified Precursors and Protein Groups.

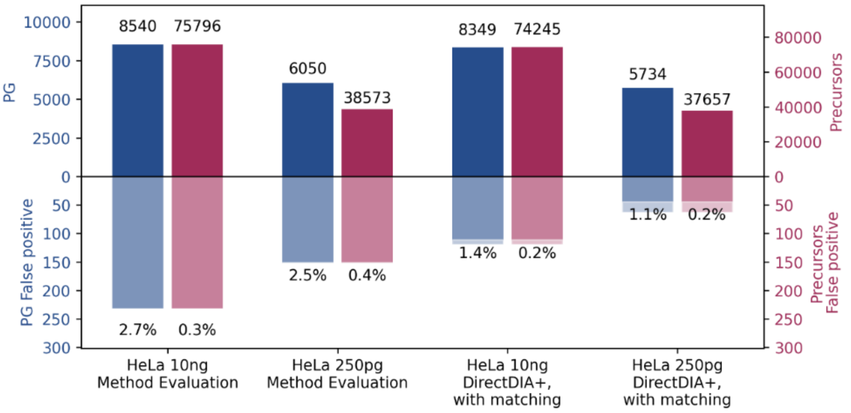

*__Supplemental Figure 2__: 250 pg or 10 ng of HeLa peptides from diluted bulk digest were injected each, reflecting the results shown in Figure 2, analyzed in the given mode using Spectronaut 18 at 1% FDR against a target (i.e. human proteome) and fake (i.e. shuffled human proteome) database. Bars indicate the total number of identified protein groups or precursors in the target database (top) and fake database (bottom) within three technical replicates.*

**Supplemental Figure 3:** Workflow benchmark, one-pot 384 well vs ProteoCHIP

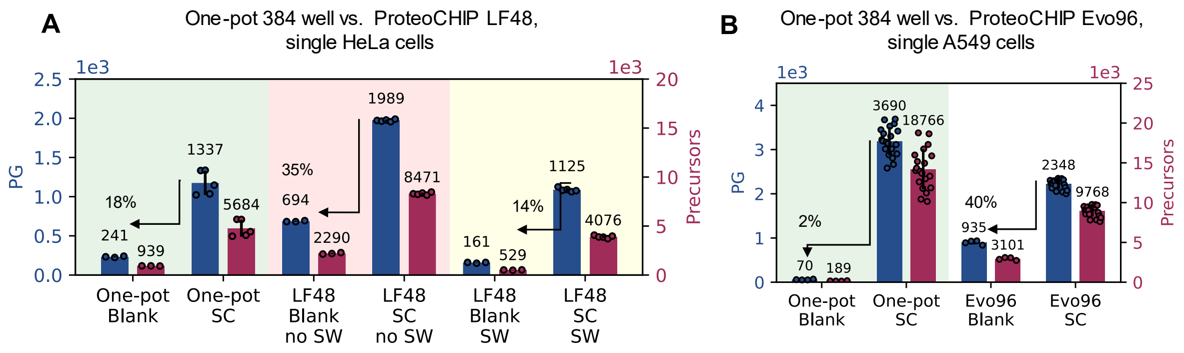

*Supplemental Figure 3: Single cells were isolated, lysed and digested using the CellenONE robot either using the previously published one-pot 384 well protocol, the LF48 or the Evo96 ProteoCHIP as indicated. Data was acquired on an Orbitrap Exploris 480 mass spectrometer at 50 SPD and data analysis was performed in directDIA+ mode using Spectronaut 18. Dots represent identified precursors or protein groups from individual replicates, bars show mean values and error bars indicate standard deviations. **A**. Comparison of one-pot 384 well (Armadillo) to the (reused) LF48 CHIP with or without an additional stringent washing (SW) to reduce background contamination. Single HeLa cells of 18-25 µm. **B**: Comparison of one-pot 384 well (Armadillo) to the (reused) Evo96 CHIP protocol. Single A549 cells of 20-30 µm.*

**Supplemental Figure 4**: Human – Yeast proteome mix to assess quantitative precision and accuracy.

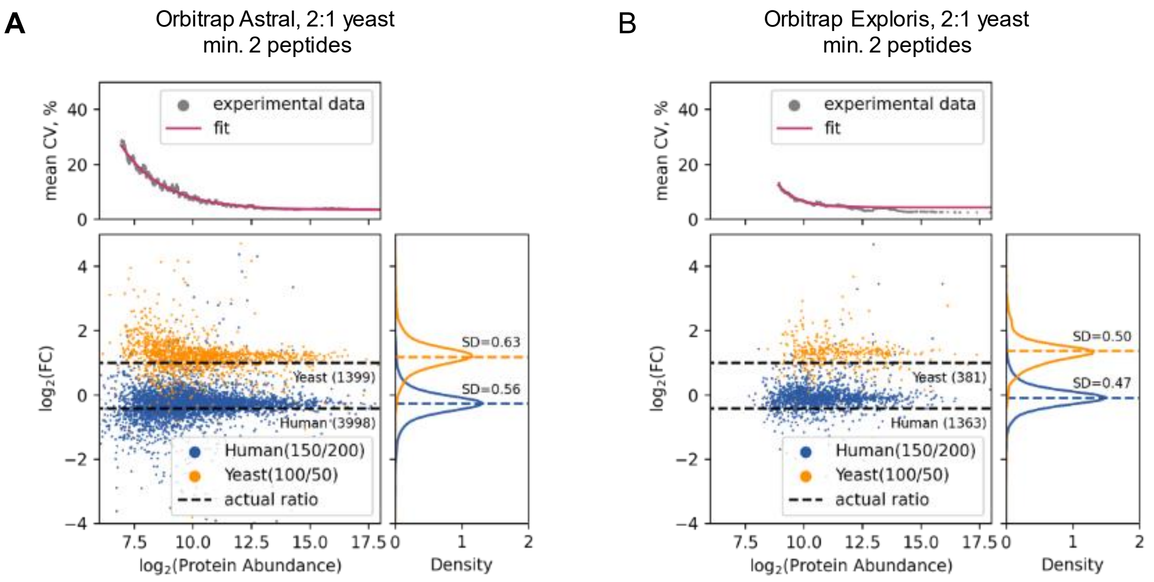

***Supplemental Figure 4**: From diluted bulk digests 250 pg two-proteome mixes consisting of either 200 pg HeLa + 50 pg yeast or 150 pg HeLa + 50 pg yeast were injected each. Peptides were separated at a throughput of 50 SPD. Data was recorded in DIA mode using optimal but not same settings for the Orbitrap Astral and Orbitrap Exploris 480 (see methods section) and analyzed using direct DIA+ in Spectronaut 18 at 1% FDR. Quantification was done on MS1 level. The same data as in Figure 5 is shown but filtered for proteins of which at least 2 peptides were quantified. Dots within the Bland–Altman plots (bottom) represent proteins with given log2 average abundance and log2 fold change of abundance across both proteome mixes. Density plots (right)depict distribution of measured log2 fold changes and CV diagrams (top) show the local CV of 100 proteins quantified with a rolling window over the entire abundance range, n=3. A: Data obtained from the Orbitrap Astral MS, B: or Orbitrap Exploris 480 MS.*

**Supplemental Figure 5**: Human – Yeast proteome mix to assess quantitative precision and accuracy, 5:1 and 10:1 yeast proteome ratio.

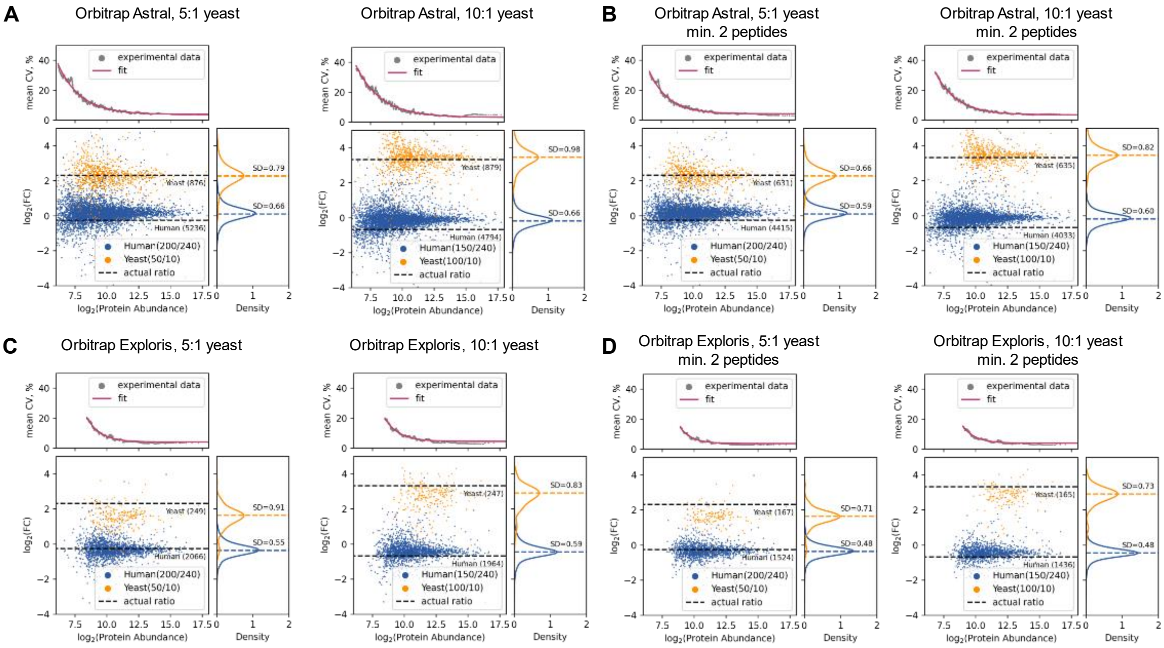

*Supplemental Figure 5: From diluted bulk digests 250 pg two-proteome mixes consisting of either 150 pg HeLa + 100 pg yeast, 200 pg HeLa + 50 pg yeast or 240 pg HeLa +10 pg yeast were injected each. Peptides were separated at a throughput of 50 SPD. Data was recorded in DIA mode using optimal but not same settings for the Orbitrap Astral and Orbitrap Exploris 480 (see methods section) and analyzed using direct DIA+ in Spectronaut 18 at 1% FDR. Quantification was done on MS1 level. Dots within the Bland–Altman plots (bottom) represent proteins with given log2 average abundance and log2 fold change of abundance across both proteome mixes. Density plots (right)depict distribution of measured log2 fold changes and CV diagrams (top) show the local CV of 100 proteins quantified with a rolling window over the entire abundance range, n=3. **A**: Data obtained from the Orbitrap Astral MS, **B**: or Orbitrap Exploris 480 MS showing all quantified proteins. **B & D**: Data for Orbitrap Astral (B) and Orbitrap Exploris 480 (D) filtered for proteins of which at least two peptides were quantified.*

**Supplemental Figure 6:** Distribution of protein abundance in A549 cells correlates to cell cycle and cell size.

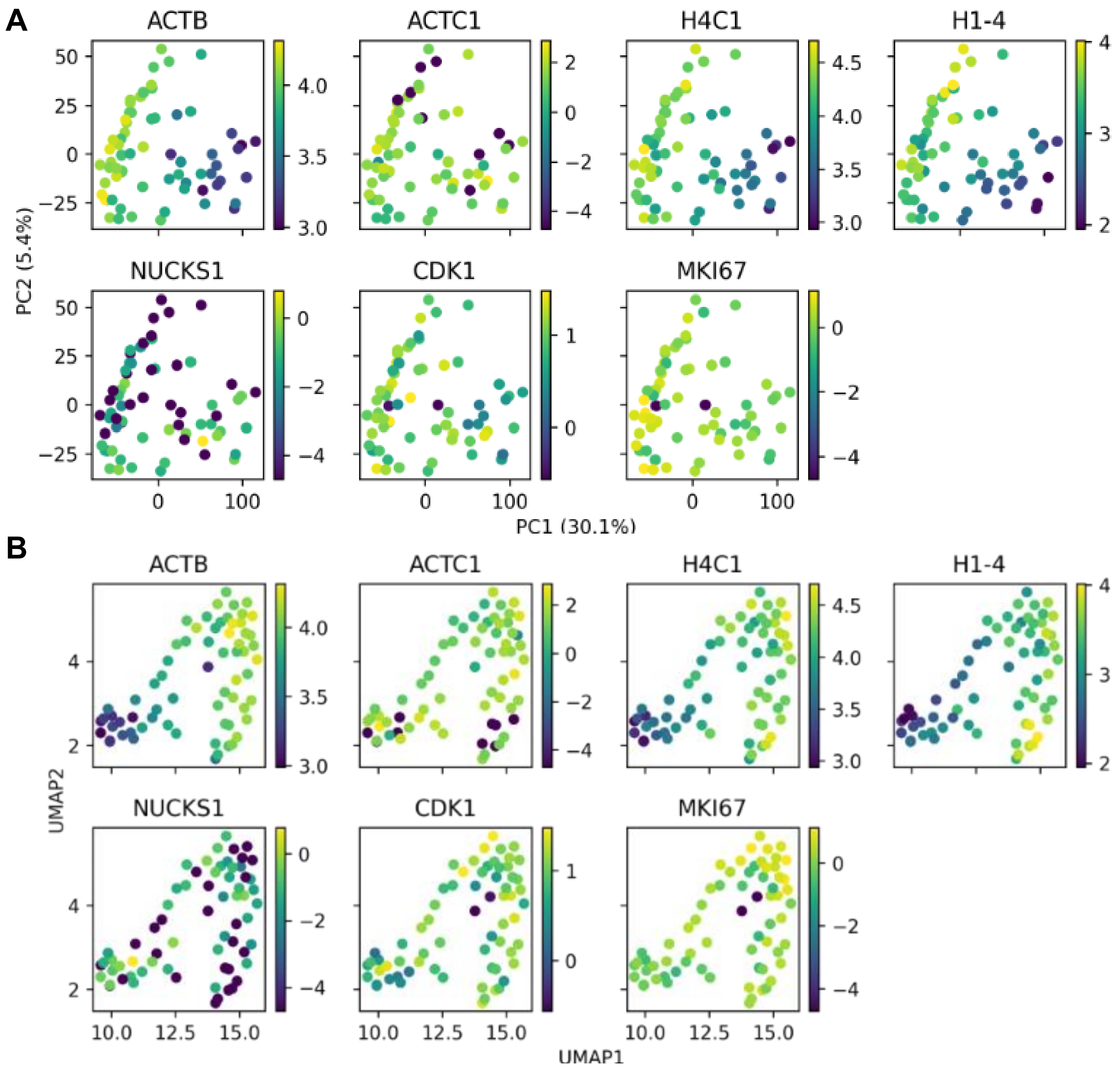

***Supplemental Figure 6**: Individual A549 cells were prepared in a 384 well plate using the CellenONE. Digested cells were analyzed at 50 SPD using the previously optimized settings for LC and MS on the Orbitrap Astral MS. The color scale bars depict the log2 protein abundance. **A**: PCA and **B**: UMAP based on protein quantities with each dot representing a cell and colors reflecting the relative abundance of the protein given.*

**Supplemental Figure 7**: Distribution of protein abundance in TE vs hPSC cells on a PCA plot.

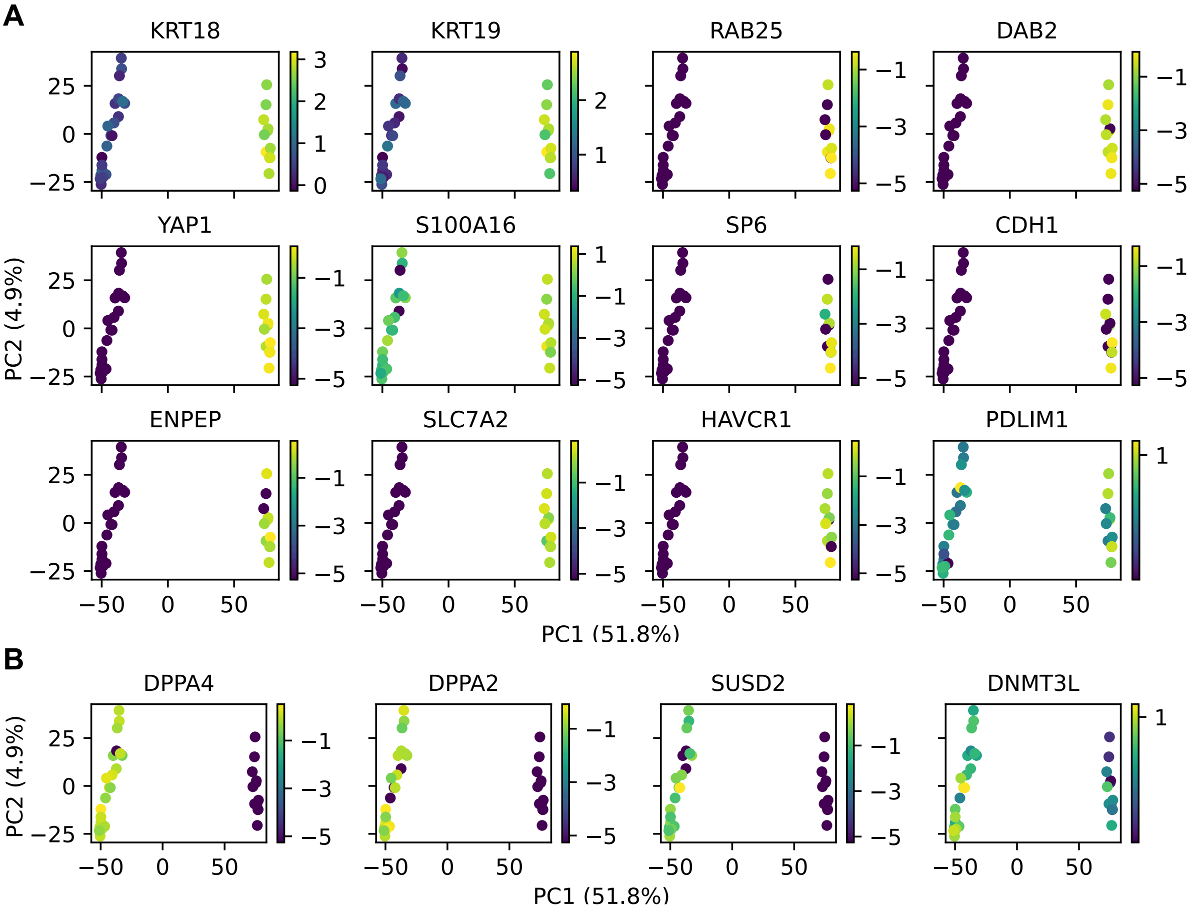

*Supplemental Figure 7: Individual hPSC and TE cells were prepared in a 384-well plate using the FACS. Digested cells were analyzed at 50 SPD using the previously optimized settings for LC and MS on the Orbitrap Astral MS. PCA on protein quantities with each dot representing a cell and colors reflecting the relative abundance of the protein given. The color bar corresponds to log₂-transformed protein abundance. Selected markers of TE cells* ***(A)*** *and hPSC* ***(B).***

**Supplemental Figure 8:** GO analysis of upregulated proteins in TE cells with highlighted selected biological pathways.

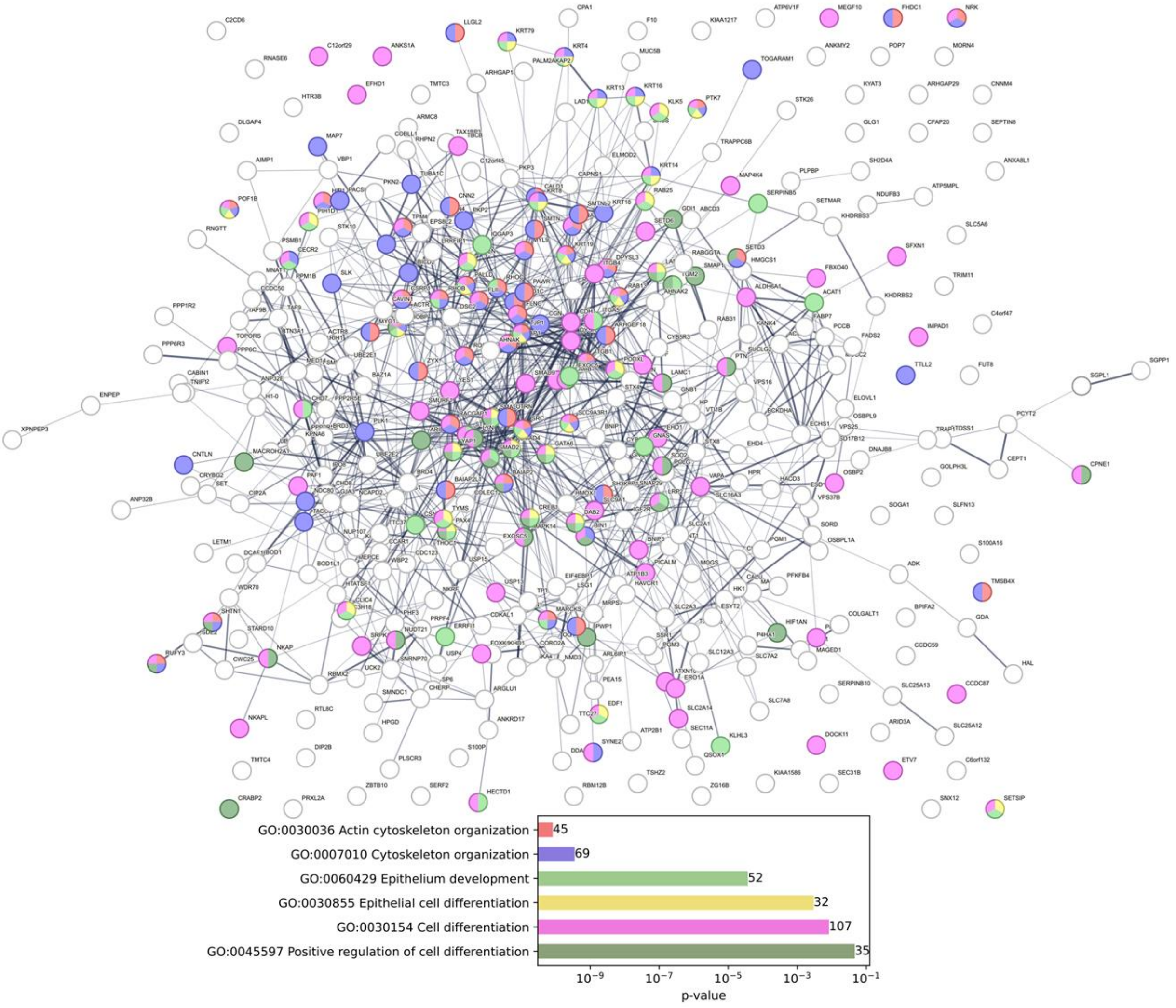

*Supplemental Figure 8: GO analysis was performed using String-db.org. At the bottom: for highlighted biological pathways p-values corrected for multiple testing within each category using the Benjamini–Hochberg procedure are shown. Number next to each bar corresponds to differentially expressed proteins found in this study for a pathway.*

**Supplemental Figure 9:** Base-peak chromatograms with and without FAIMS

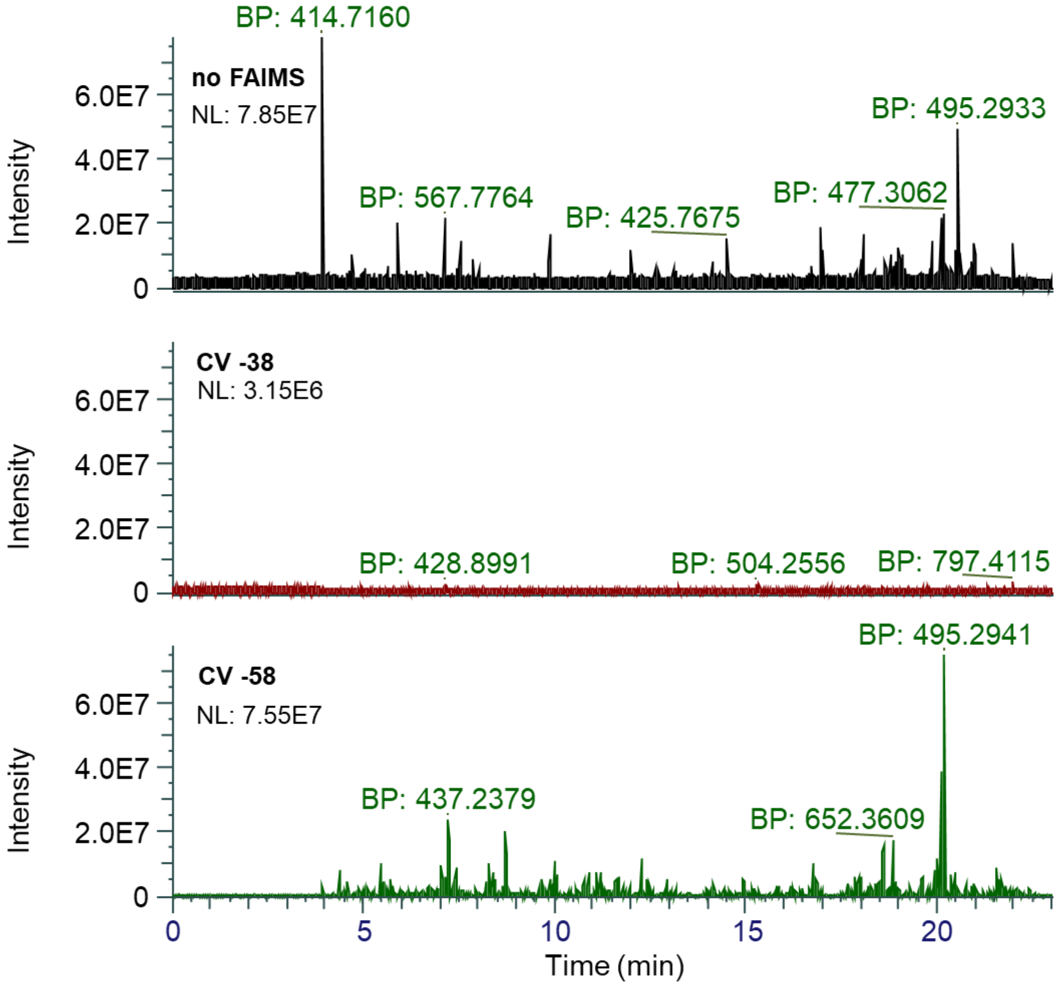

*Supplemental Figure 9: Base-peak chromatograms from representative replicate measurements from the data shown in Figure 4. 250 pg of HeLa peptides from diluted bulk digest were injected each. Peptides were separated at a throughput of 50 SPD. Data was recorded with or without the FAIMS unit attached and at the given compensation voltage.*

**Supplemental Figure 10:** Intersection of identified peptides in runs with and without FAIMS Pro interface.

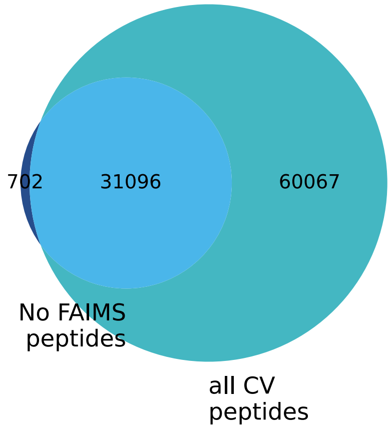

*Supplemental Figure 10: Venn diagram showing the intersection of identified peptides in all acquired runs with and without FAIMS Pro interface. Rus were acquired from 250 pg of HeLa peptides from diluted bulk digest. Peptides were separated at a throughput of 50 SPD. Data was recorded with or without the FAIMS Pro Duo interface unit attached and at the compensation voltage in range -38 V to -88 V with 10 V step.*
